# Parental Allele-Specific Protein Expression in Single Cells *In Vivo*

**DOI:** 10.1101/413849

**Authors:** Chiu-An Lo, Brian E. Chen

## Abstract

Allelic expression from each parent-of-origin is important as a backup and to ensure that enough protein products of a gene are produced. Thus far, it is not known how each cell throughout a tissue differs in parental allele expression at the level of protein synthesis. Here, we measure the expression of the Ribosomal protein L13a (Rpl13a) from both parental alleles simultaneously in single cells in the living animal. We use genome-edited *Drosophila* that have a quantitative reporter of protein synthesis inserted into the endogenous *Rpl13a* locus. We find that individual cells can have large (>10-fold) differences in protein expression between the two parental alleles. Cells can produce protein from only one allele oftentimes, and time-lapse imaging of protein production from each parental allele in each cell showed that the imbalance in expression from one parental allele over the other can invert over time.

**One sentence summary:** Parental allele-specific protein expression varies widely across cells and over time.

**Highlights:**

1. We used genome editing to insert a quantifiable protein translation reporter into the endogenous *Ribosomal protein L13a* gene and thus track the protein expression of both parental alleles simultaneously in every single cell in the awake animal.
2. Cells can have a large difference in protein expression for one parental allele over the other, and this can invert over time, and can occur in clusters of cells within a tissue.
3. We demonstrate the highly variable nature of heterozygous and homozygous definitions across single cells, and over time.
4. Our study demonstrates a new paradigm that can be used to examine inherited epigenetic control of expression from a specific parent-of-origin allele across single clonal cells from a common progenitor *in vivo*.

## Introduction

Protein synthesis is the limiting step in generating the final molecular outcome of DNA. We sought to compare protein expression among different individual cells between tissues, between each parental allele, and over time. However, it has not been possible to measure allele-specific protein expression dynamics in single cells. Thus, in order to track and quantify the dynamics of protein synthesis in single cells in animals, we used the Protein Quantitation Ratioing (PQR) technique (Figure 1A) (Lo et al. 2015). Using CRISPR-Cas9 genome editing of *Drosophila melanogaster*, we inserted a *PQR* DNA construct with a red fluorescent protein (RFP) reporter at the end of the coding sequence for the *Ribosomal protein L13a* (*Rpl13a*) gene (Lo et al. 2015) (Figure 1A). This animal produces one molecule of RFP for every one molecule of Rpl13a protein that is made during protein synthesis. Therefore, the red fluorescence intensity in a cell is proportional to the Rpl13a concentration and can be used to track and measure Rpl13a dynamics over time in every cell in the animal. We chose Rpl13a because, as a ribosomal subunit, it is itself involved in protein translation and has a long turnover rate similar to fluorescent proteins (Mazumder et al. 2003; Chaudhuri et al. 2007; Kapasi et al. 2007; Boisvert et al. 2012; Jia et al. 2012; Lo et al. 2015). Rpl13a is expressed in all cells at moderately high levels with several hundreds of mRNA copies per cell (Lo et al. 2015). *Rpl13a* mRNA expression dynamics in the brain and in the whole organism are not circadian driven in flies and mice (Keegan et al. 2007; Hughes et al. 2010; Hughes et al. 2012). Rpl13a is also frequently used as a control “housekeeping” gene in quantitative DNA measurements because of its resistance to external effects and stability (Mane et al. 2008).

**Figure 1.**
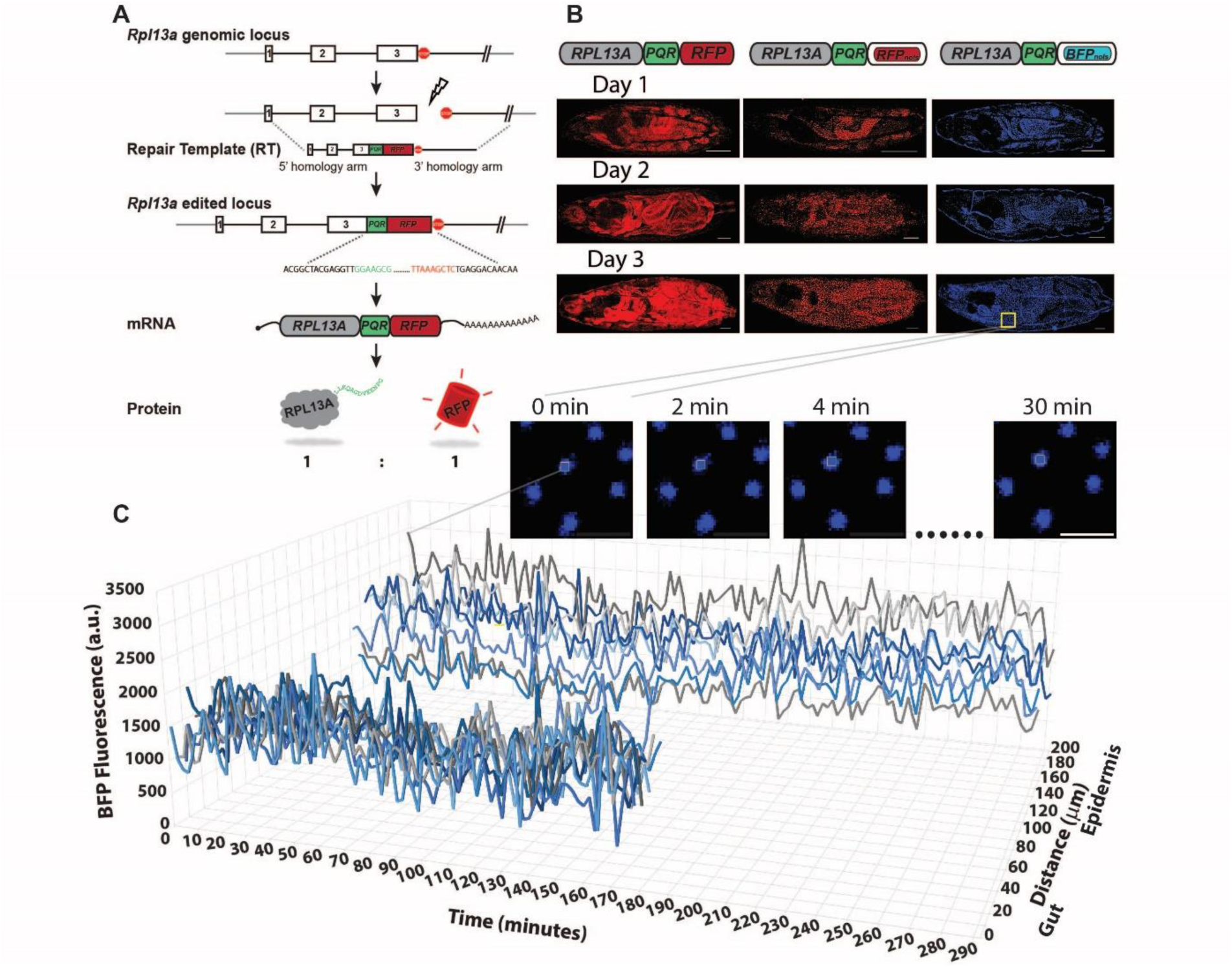
Quantification of Rpl13a protein production in single cells in the living animal. (A) Insertion of a *Protein Quantitation Reporter* (*PQR*) into the endogenous *Rpl13a* gene can quantify protein amounts. The *PQR* is inserted before the final stop codon using CRISPR-Cas9 genome editing (lightning bolt). The break is repaired using homologous recombination of an exogenous repair template containing the *PQR* and appropriate homology arms. Colored nucleotide sequences represent genomic sequencing results of a *PQR-RFP* insertion into *Drosophila melanogaster*. During protein synthesis the *PQR* produces a stoichiometric ratio of red fluorescent protein (RFP) and Rpl13a protein. (B) Rpl13a increases expression across the animal over days. Three different lines of genome-edited *Drosophila melanogaster* were created, Rpl13a-PQR-RFP, Rpl13a-PQR-RFP_nols_ and Rpl13a-PQR-BFP_nols_. A nucleolar localization signal was used to sequester the RFP or blue fluorescent protein into the nucleus (RFP_nols_ or BFP_nols_, respectively) to visualize individual cells.Fluorescence intensity in each nucleus is used to measure Rpl13a protein expression. Scale bars, 100 µm (main images), 10 µm (inset). (C) Rpl13a protein expression changes can be measured over hours. Homozygous Rpl13a-PQR-BFP_nols_ larvae were restrained and imaged every 2 minutes for 5 hours. Blue fluorescence intensities from 8 cells in the epidermis and 10 cells from the gut are shown.

## Results

### Measuring protein synthesis dynamics in single cells in the awake animal

Inserting *PQR-RFP* into the endogenous *Rpl13a* locus resulted in an entirely red fluorescent animal (Figure 1B). These *Drosophila* were easily identified by their strong red fluorescence everywhere, which increased globally as the animal developed (Figure 1B). Although individual cells were not distinguishable in these animals, we used these animals to verify that global Rpl13a is arrhythmic over hourly timescales. To distinguish individual cells, we created two more genome-edited *Drosophila* PQR lines that sequestered RFP or blue fluorescent protein (BFP) into the nucleus, Rpl13a-PQR-RFP_nols_ and Rpl13a-PQR-BFP_nols_ animals, respectively (Figure 1B). Nuclear localization signals tend to result in fluorescent proteins leaking into the cytosol. We added a nucleolar localization signal (Tsai et al. 2008) to RFP (RFP_nols_) and BFP (BFP_nols_) because nucleolar localized fluorescent proteins spread only into the nucleus (Lo et al. 2015).

We tracked endogenous Rpl13a protein production in single cells *in vivo* by measuring red and blue fluorescence intensities in the nucleus over time scales of seconds to days (**Methods**). To avoid the effects of anesthesia on protein translation, we imaged awake animals immobilized using a suction microfluidics chamber (Mishra et al. 2014). Images were acquired at varying time intervals to verify that the time course of fluorescence signals were not due to changes in animal positioning, movement, imaging depth, photobleaching, or changes in the nucleus and nucleolus (Figure 1C**, Methods**). At the larval stages after *Drosophila* embryos hatch, mitosis ceases and nearly all cells are fully differentiated and are simply expanding rapidly in size without dividing (Smith and Orr-Weaver 1991).

### Individual cells can have remarkably imbalanced protein expression for one allele

Mothers produce the oocyte and contribute to it nutrients, proteins, and mRNA molecules, until zygotic expression of both the maternal and paternal genes turn on. We sought to examine the dynamics of protein expression between each of the parent-of-origin *Rpl13a* alleles after zygotic expression turns on. We took advantage of the two colors of the PQR lines and crossed them to create transheterozygous *Rpl13a-PQR-RFP_nols_ / Rpl13a-PQR-BFP_nols_* animals, from Rpl13a-PQR-RFP_nols_ mothers and Rpl13a-PQR-BFP_nols_ fathers, or vice versa (Figure 2), to track expression of each allele. We verified that oocytes from Rpl13a-PQR-RFP_nols_ mothers had only red (i.e., maternal expression) within cells (Figure 2A), until the dark period when all maternal expression is degraded during the maternal to zygotic transition (Tadros and Lipshitz 2005; Schier 2007; Baroux et al. 2008). Upon zygotic (i.e., genomic) expression of Rpl13a protein, we found that both parental alleles were expressed at the same time across the organism (Figure 2A).

**Figure 2.**
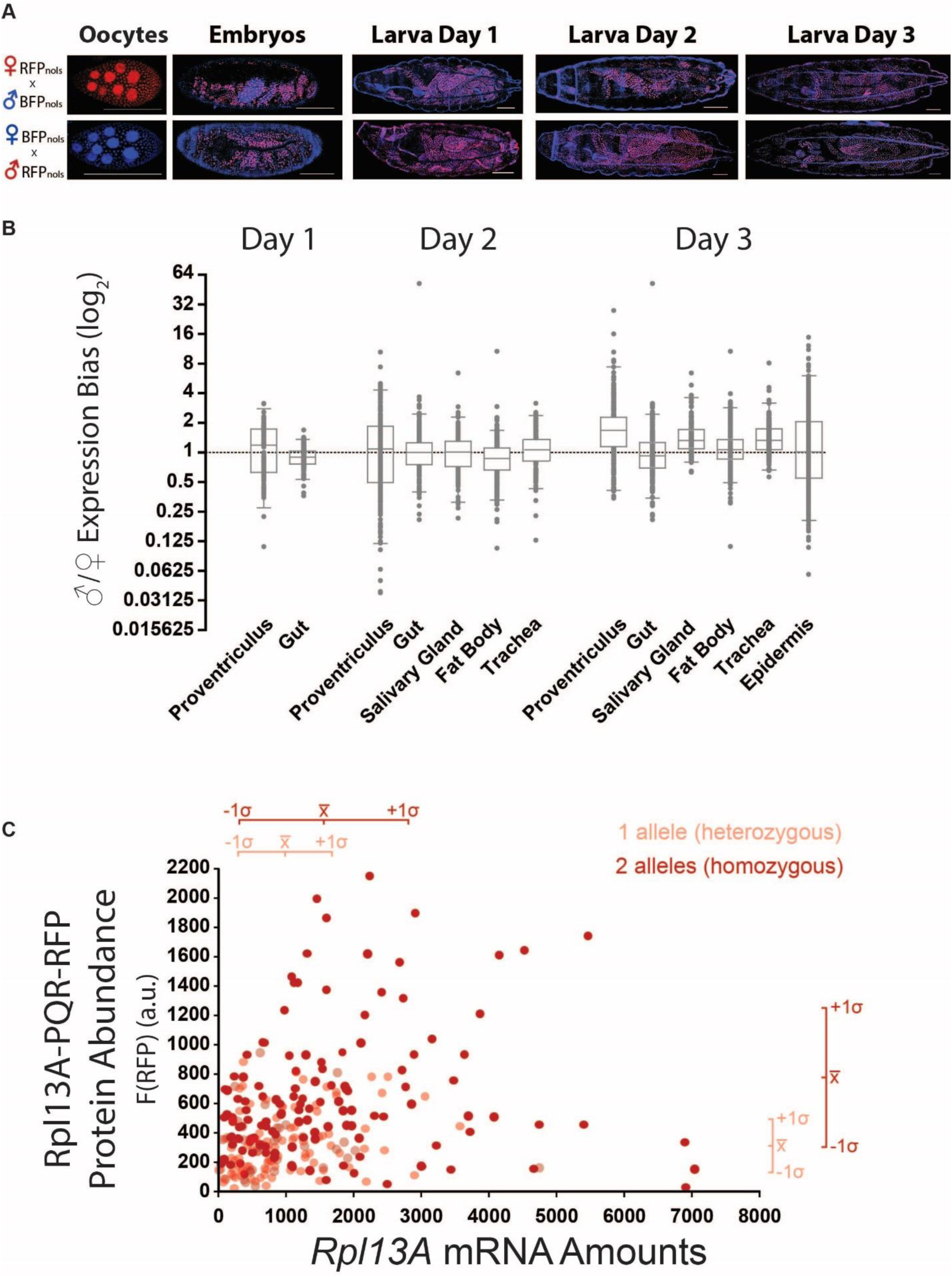
Large differences between maternal versus paternal allele protein expression can occur in single cells. (A) Rpl13a-PQR-RFP_nols_ and Rpl13a-PQR-BFP_nols_ parents produced transheterozygous progeny whose specific parent-of-origin Rpl13a allele can be tracked using color. Maternal expression of Rpl13a was only detected in oocytes, where it is expressed at high levels and then decreases rapidly in embryos. Rpl13a expression is only detectable again in late embryos, simultaneously from both the maternal and paternal alleles. Most cells express both alleles equally, relative to the blue and red fluorescence intensities, but clusters of cells within the same tissue had similar allelic biases. Scale bars, 100 µm. (B) The ratio of the paternal to maternal Rpl13a protein expression was measured in single cells across different tissues and from both combinations of Rpl13a-PQR-RFP_nols_ and Rpl13a-PQR-BFP_nols_ parents. Box plots represent the median and 1^st^ and 3^rd^ quartiles of the data and error bars are 2.5 and 97.5 percentiles. (C) The relationships between DNA allele number, RNA, and protein numbers. Single cells were isolated from heterozygous and homozygous Rpl13a-PQR-RFP_nols_ animals and their fluorescence intensities and mRNA amounts were measured. Despite the weak correlation between mRNA and protein levels, the difference between having one copy of DNA expressed versus having two copies of the allele is evident from the population data. Stochastically, homozygous animals are able to reach mRNA and protein levels greater than a single allele is, but mRNA and protein levels from two alleles can often be less than that of a single allele.

What percentage of cells express both the mother’s and father’s allele at any given time? Imaging transheterozygous larvae at different days, we found that all cells had both red and blue nuclei, as might be expected for a gene expressed at fairly high levels. The majority of cells had similar fluorescence intensities of RFP and BFP (Figure 2A). We wanted to know whether there were equivalent levels of mRNA between the parental alleles which might roughly correlate to the similar levels of red and blue fluorescence observed, even though we and others have shown that mRNA amounts correlate poorly with protein quantity, including for *Rpl13a* (Schwanhausser et al. 2011; Vogel and Marcotte 2012; Lo et al. 2015). We performed qPCR on single cells isolated from transheterozygous *Rpl13a-PQR-RFP_nols_ / Rpl13a-PQR-BFP_nols_* animals and found 941 ± 1066 mRNA molecules from the maternal allele and 943 ± 878 mRNA molecules from the paternal allele (*n* = 53 cells, 32 animals, 12 from PQR-RFP_nols_ fathers).

These results show that the broadly similar levels of red versus blue fluorescence intensities across the animal do correspond with broadly similar levels of mRNA expression from the respective alleles. We next sought to measure the relative differences between the paternal versus maternal Rpl13a protein expression in individual cells, by adjusting the fluorescence imaging acquisition to be equivalent values between red and blue fluorescence for most cells in the animal. Using the ratio between the red to blue PQR fluorescence expression in each nucleus, we measured the distribution of relative contributions of each parental allele in cells in the proventriculus, gut, salivary gland, fat body, trachea, and epidermis. Most cells expressed both alleles relatively equally, but we frequently observed an imbalance for one parental allele over the other in single cells (*n* = 17 animals, 7 from PQR-RFP_nols_ fathers and 10 from PQR-BFP_nols_ fathers). Occasionally we observed clusters of cells that all had the same imbalanced expression for one allele (Figure 2A), indicating a clonally inherited initial bias set by early common progenitor cells, but we never observed a complete silencing or systematic bias across the animal for a parental allele (Wang et al. 2008; Gregg et al. 2010a; Gregg et al. 2010b; DeVeale et al. 2012; Xie et al. 2012; Crowley et al. 2015). Some cells express very little of one allele leading to differences as large as 64-fold between the two alleles (Figure 2B). Thus, it is important to note that at the protein level each parental allele in an animal will not necessarily be uniformly expressed in all cells.

Does a heterozygous animal express half as much RNA or protein as a homozygous animal? Our single cell qPCR results show that not every cell transcribes an equal amount of each parental allele, and this is also the case at the protein level, with some cells barely expressing an allele (Figure 2B). In other words, if the choice to express either parental allele is stochastic, and the correlation between RNA and protein is poor, then many cells in a heterozygous animal might not necessarily be heterozygous at the RNA nor protein levels. We sought to visualize this relationship between DNA to RNA to protein distributions (Figure 2C). We isolated 129 single cells from 34 transheterozygous Rpl13a-PQR-RFPnols / Rpl13a-PQR-BFPnols animals and 138 cells from 24 homozygous Rpl13a-PQR-RFPnols animals and then measured their PQR fluorescence intensities before performing single cell qPCR. The linear correlation between *Rpl13a* mRNA and protein expression was weak, with coefficients of determination, *R*^2^ values close to 0. However, the average number of *RFP* mRNA in single cells in the heterozygotes was 979 ± 776 compared to 1650 ± 1241 in the homozygotes. The RFP fluorescence intensity per heterozygous cell was 338 ± 209 compared to 738 ± 484 in the homozygotes. Thus, even with stochastic mechanisms governing the DNA transcription of each allele, RNA expression amounts, and protein synthesis, a large number of cells in an organism averages out the noise to produce more protein from two copies of DNA versus a single copy of the allele.

### Allele-specific imbalances in protein expression change over time

Does a cell maintain its ratio of parental allele expression over time? To measure the protein expression dynamics of both parental alleles, we imaged transheterozygous *Rpl13a-PQR*-*RFP_nols_ / Rpl13a-PQR-BFP_nols_* awake animals every 15 minutes over time periods of 5 hours (*n* = 8 animals, 4 from PQR-RFP_nols_ fathers; Figure 3A–C). Tracking the ratio of expression between the paternal and maternal Rpl13a allele over time from 144 cells revealed that most cells did not show more than a 4-fold bias for either parental allele (Figure 3D). The average paternal/maternal ratios in single cells over time from different tissues were 1.186 ± 0.858 (epidermis), 1.044 ± 0.709 (salivary gland), and 0.960 ± 0.419 (proventriculus) (mean ± S.D., *n* = 5 animals, 3 from PQR-RFP_nols_ fathers). Thus, over time the majority of cells express close to equal amounts between the two parental alleles, but still many cells will express a > 4-fold difference between the two alleles (Figure 2B, 3D). Interestingly, this bias can invert, as we found that 21% of cells that expressed a > 2-fold difference between the parental alleles would invert their allele preference ratio to a > 2-fold bias for the other allele (Figure 3D). This emphasizes the dynamic nature of allelic expression and shows that the heterozygosity of a single cell can dramatically change over time periods of hours.

**Figure 3.**
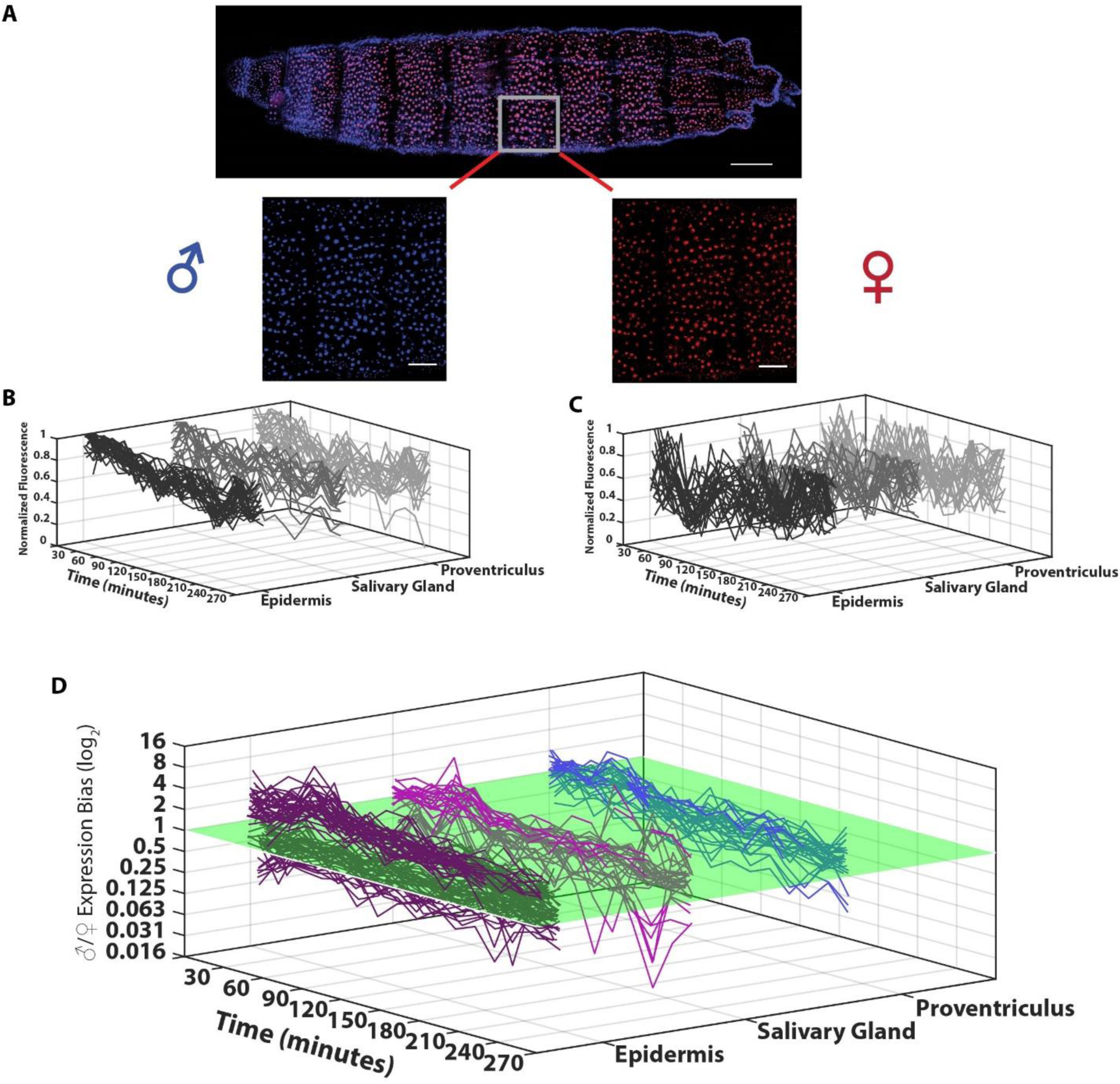
Parental allele-specific protein expression can change dramatically over time. (A–C) Each parent-of-origin allele of Rpl13a can be tracked and quantified in Rpl13a-PQR-RFP_nols_/Rpl13a-PQR-BFP_nols_ transheterozygous animals using the blue and red fluorescence intensities of the cell nucleus (**A**). Protein synthesis dynamics from paternal (**B**) and maternal (**C**) alleles from each cell in different tissues were measured at 15-minute intervals. Scale bars, 100 µm in whole animal image, 50 µm in magnified images. (D) The ratio of paternal to maternal allele expression changes over time. Each curve represents the fluorescence ratio from an individual cell. The surface plane denotes equal expression of paternal and maternal alleles. Curves crossing the surface plane represent an inversion of allele preference expression at the time. The majority of cells maintain close to equal expression of each allele over timescales of hours. However, at any given time many cells will express a > 2-fold expression preference for one parental allele, and even this bias can invert over time.

## Discussion

We used fluorescent proteins to report the protein expression of Rpl13a. After protein synthesis of Rpl13a and the PQR fluorescent protein reporter, both proteins must fold and mature to function. It is not known how long the Rpl13a protein takes to function after its synthesis, but it is similar in size to the red and blue fluorescent proteins used in our experiments, which we estimate to take one hour to fluoresce after synthesis at 25°C (Balleza et al. 2018). Therefore, we analyzed the ratios of fluorescence intensities between cells and between alleles rather than quantifying amounts of functional Rpl13a.

Although protein activity is the ultimate executor of gene expression, RNA levels are frequently used as a proxy, and so knowing the correlation between a gene’s mRNA and protein products is important to many biologists. For example, up to half of proteins with circadian rhythms do not have rhythmic mRNAs (Reddy et al. 2006; Robles et al. 2014). Protein abundance is critically important in some genes, where the loss of expression of one copy of the gene can cause diseases and disorders collectively called haploinsufficiencies. Our results examining allelic expression at the RNA and protein levels demonstrate that heterozygous and homozygous definitions vary from cell to cell and over time. For example, this asymmetry could allow one parental allele with a disease mutation to be the one solely expressed during transcription or spatially during local translation in a neuron’s dendrite. Overall within a tissue, the protein expression between two alleles balances out (Figure 2B), but still, close to 50% of cells in an animal can have a > 2-fold difference between the two alleles. Furthermore, these biases can change over time in single cells (Figure 3D), further highlighting the fluid definition of allelic expression. Despite the noise in mRNA and protein production, the underlying structure that is set by the integer copy numbers of DNA (i.e., one allele expression versus two allele expression) is clear and detectable with even a moderate number of cells (Figure 2C). These cellular and temporal differences in allelic expression have important consequences for understanding diseases caused by haploinsufficiency and copy number variations. Our study demonstrates a new paradigm to examine parental allele-specific expression at the protein level in single cells over time in the living animal. This new approach can be useful in examining genes subject to imprinting where only a specific parent-of-origin allele is expressed due to the presence of inherited epigenetic controllers. Tracking the inheritance of epigenetic signatures that control allelic expression across multiple cell divisions of individual cells can be performed in real time *in vivo* using this approach.

## Experimental Procedures

### Preparation of sgRNA Expression vectors

Different single guide RNAs (sgRNAs) were designed in a 20 base pair DNA oligonucleotide format to guide the Cas9 nuclease to the end of the coding region of *Drosophila Rpl13a*. They were optimized using the on-line Optimal Target Finder algorithm (http://tools.flycrispr.molbio.wisc.edu/targetFinder). Individual sgRNAs were synthesized and cloned into the backbone of the *pCFD3: U6:3-gRNA* vector (Addgene #49410) (Port et al. 2014). Each sgRNAs was co-transfected along with a Cas9 expressing plasmid, pBS-Hsp70-Cas9 (Addgene #46294), and a circular *Rpl13a* repair template into cultured *Drosophila S2* cells using TransIT Insect Transfection Reagent (Mirus Bio LLC, Madison, WI, USA). Gene editing efficiency of the different sgRNAs was evaluated five days after transfection by cellular fluorescence and genotyping using a primer outside of the homology arm and the other inside the inserted *PQR* sequence.

### Construction of *PQR* Repair Template

The genomic sequences of *Drosophila Ribosomal protein L13A* were identified using GeneDig (https://genedig.org) (Suciu et al. 2015) and then PCR amplified from Canton-S fly genomic DNA lysates to construct the 5’ and 3’ homology arms of 1.4 kilobase each. The 5’ homology arm did not include the endogenous *Rpl13a* promoter, to prevent the expression of the transgene until the in-frame genomic integration at the correct locus. The three different *Protein Quantitation Reporters* with a fluorescent protein with and without a nucleolar localization signal were inserted between the 5’ and 3’ homology arms to generate three different *Rpl13a*-specific repair templates as described (Lo et al. 2015).

### Generation of sgRNA-expressing transgenic fly

The optimal sgRNA in *pCFD3: U6:3-gRNA* vector was microinjected into embryos (BestGene, Inc) to create a transgenic fly constitutively expressing the *Rpl13a*-specific sgRNA. This sgRNA fly was then crossed to a *nos-Cas9* fly (*yw*; *attP40 {nos-Cas9}*/*CyO*) to restrict CRISPR-Cas9 activity to minimize lethality (Port et al. 2014). Circular *Rpl13a* repair template was then microinjected into the embryos containing both active Cas9 and sgRNA (BestGene, Inc). Surviving flies were intercrossed with one another and the resulting offspring were screened for red or blue fluorescence signals at wandering 3^rd^ instar larval stage. RFP or BFP positive larvae were collected and out-crossed to the Canton-S strain to remove the Cas9 and sgRNA genes. The integrated *PQR-RFP, PQR-RFP_nols_*, or *PQR-BFP_nols_* at the *Rpl13a* locus at the 3^rd^ chromosome was balanced by crossing to a TM3 balancer line. All three of the genome-edited fly lines (Rpl13a-PQR-RFP, Rpl13a-PQR-RFP_nols_ and Rpl13a-PQR-BFP_nols_) were verified by genotyping and sequencing.

### Fly husbandry

Flies of different genotypes were reared in a 25°C humidity controlled incubator with a 12:12 hour light/dark cycle. Embryos were collected for 4 hours (stage 12-16) on apple juice agar plates and the hatched larvae were imaged at the next day or 2 or 3 days later. Homozygous and transheterozygous embryos or larvae were collected for imaging. Oocytes from transheterozygous females were used to confirm maternal expression of Rpl13a, which was expressed strongly, but then decreased rapidly in embryos.

EHMT null *(EHMT^DD1^;; Rpl13a-PQR-RFP_nols_ / Rpl13a-PQR-BFP_nols_* and *EHMTDD2;; Rpl13a-PQR-RFP_nols_ / Rpl13a-PQR-BFP_nols_)* larvae were obtained by crossing *w;; Rpl13a-PQR-RFP_nols_/TM3* to *EHMT^null^* imprecise p-element excision flies (gifts from Dr. Annette Schenck). The control EHMT fly line used was the precise excision of the same genetic background (Kramer et al. 2011). Larvae were used at 3 days old for experiments. UNC0638 containing apple agar plates were used to collect embryos and used for the larvae food source. These larvae were used at 3 days old for experiments.

### Quantitative Real-Time PCR

Single cell imaging and subsequent qPCR was performed as previously described (Lo et al. 2015). Individual cells were imaged in drops of culture media on Teflon-coated glass slides before extraction and purification of total RNA using the TRIzol reagent (Life Technologies). Total RNA was reverse-transcribed with gene-specific primers (2µM final concentration) using Superscript III reverse-polymerase (Life Technologies). This cDNA template was used for real-time PCR using the TaqMan Fast Advanced Mastermix (Life Technologies). Real-time PCR amplification was detected using the StepOnePlus Real-Time PCR System (Applied Biosystems) and cycle quantification values were calculated using the StepOne software. Experiments were performed in two to three experimental replicates with two technical replicates. Absolute quantification was determined using standard curves generated with synthesized oligo standards containing the *Rpl13a* target. Primers specific for *Rpl13a-PQR-RFP* and *Rpl13a-PQR-BFP* were less efficient (86%) than primers for wildtype *Rpl13a*, which can result in less accurate absolute RNA quantifications. Primers and double-quenched 5’-FAM/ZEN/IowaBlackFQ-3’ probes were purchased from Integrated DNA Technologies (Coralville, IA).

### Image Acquisition

Fluorescence images were taken using an Olympus laser scanning confocal microscope FV1000 at 800 × 800 pixels with a 20× or 40× oil objective, N.A. 0.85 and 1.30 corresponding to a 636 × 636 µm and 317 × 317 µm field of view, respectively, or a custom built 2-photon laser scanning microscope at 512 × 512 pixels with a 40× water objective, 1.0 N.A. corresponding to an approximately 310 × 310 µm field of view. Fluorescence emission was detected by photomultiplier tubes. All image acquisition parameters were fixed during imaging sessions for excitation intensity, exposure time, gain, and voltages. Animals were imaged at late stage embryos (>16 hours after egg laying), and three different developmental stages (1, 2, and 3 days after hatching) of the transparent larvae. For long term time-lapse imaging, larvae were immobilized and accommodated in a microfluidic chamber (larva chip) over the course of imaging (Mishra et al. 2014). Larvae were coated with Halocarbon 700 oil to avoid dehydration. Images were acquired between every 60 seconds, 2, 4, 5, or 15 minutes for 3 to 7 hours total duration to verify that the time constant for fluorescence changes (see below) were not due to animal positioning, movement, imaging depth, or photobleaching, regardless of sampling frequency.

### Image Analysis

Average fluorescence pixel intensities were measured in one region of interest (of 2 × 2, 4 × 4, or 5 × 5 pixels for epidermal, fat body, or gut cells, respectively) that covered 70% of the nuclei using ImageJ as described (Lo et al. 2015). Fluorescence pixel intensities were background subtracted and presented in arbitrary units. No bleed-through was detected between red and blue fluorescence channels. Images in figures were adjusted for contrast and brightness for presentation. To verify that changes in fluorescence intensity over time were not due to changes in nucleolar and nucleus size and shape, we imaged Rpl13a-PQR-RFP animals where the RFP fluorescence is within the cytosol (Figure 1B). In initial experiments we crossed Rpl13a-PQR-RFP with Rpl13a-PQR-BFP_nols_ animals to use the blue nucleus as a marker for individual cells, but the RFP was found to dimerize with the BFP and accumulate in the nucleus, as we have observed previously in mammalian cells. In Rpl13a-PQR-RFP animals, we analyzed cytoplasmic RFP fluorescence dynamics in putative single cells using one region of interest of 6 × 6 pixels. The fluorescence dynamics were not significantly different from animals with fluorescence contained within the nucleus, with an average time constant of τ = 8.3 ± 7.8 minutes from cytoplasmic fluorescence (mean ± S.D., *n* = 36 cells, 3 animals) compared to 8.5 ± 8.1 minutes from the nuclear fluorescence.

### Statistical Analysis

All statistical analyses were performed using custom-written programs in MatLab (MathWorks, Natick, MA). RNA and protein simulations comparing heterozygous and homozygous alleles were performed using custom-written programs (available on request) in MatLab.

## Acknowledgments

The authors thank Ibrahim Kays, Farida Emran, Júnia Vieira dos Santos, Isabela Fabri Karam, Leah Dawson, and ChenHui Zhao for assistance with experiments and analysis. This work was supported by grants (to B.E.C.) from the Natural Sciences and Engineering Research Council of Canada, and the Canadian Institutes of Health Research (148882).

## Author Contributions

B.E.C. designed the experiments and supervised the project. C.L. and B.E.C. performed experiments and analyzed the data. C.L. and B.E.C. wrote the manuscript.

No competing financial interests to declare.

Correspondence and requests for materials should be addressed to

